# An Investigation of Detection Biases in the Unattended Periphery During Simulated Driving

**DOI:** 10.1101/257741

**Authors:** Musen Kingsley Li, Hakwan Lau, Brian Odegaard

## Abstract

While people often think they veridically perceive much of the visual surround, recent findings indicate that when asked to detect targets such as gratings embedded in visual noise, observers make more false alarms in the unattended periphery. Do these results from psychophysics studies generalize to naturalistic settings? We used a modern game engine to create a simulated driving environment where participants (as drivers) had to make judgments about the colors of pedestrians’ clothing in the periphery. Confirming our hypothesis based on previous psychophysics studies, we found that subjects showed liberal biases for unattended locations when detecting specific colors of pedestrians’ clothing. A second experiment showed that this finding was not simply due to a confirmation bias in decision-making when subjects were uncertain. Together, these results support the idea that in everyday visual experience, there is subjective inflation of experienced detail in the periphery, which may happen at the decisional level.

## Introduction

How do we make perceptual decisions in the visual periphery? Subjectively, it may seem that we perceive the unattended visual periphery in precise detail, and there is some evidence supporting this claim (Block, 2007, 2011; Lamme, 2004, 2006; Sperling, 1960). However, inattentional blindness (Simons & Chabris, 1999) and change blindness (Grimes, 1996; Rensink, O’Regan, & Clark, 1997) experiments show that alterations to unattended elements in natural scenes are often unnoticed. Additionally, recent experiments reveal that coarse summary statistical properties can be perceived and remembered for unattended items, but individual details are lost (Ward, Bear, & Scholl, 2016). Profound deficits are also revealed when subjects are asked simple questions about color perception in the periphery (see Box 2, Cohen, Dennett, & Kanwisher, 2016). Thus, while perceptual deficits in the periphery are evident, a precise characterization of this deficit remains elusive, and overall the idea of impaired peripheral perception seems at odds with the subjective impression that we perceive the visual world in relatively uniform detail.

Previous work has addressed this question within the framework of Signal Detection Theory. Results show that observers tend to use liberal detection biases when evaluating peripheral or unattended stimuli, such as grating patterns embedded in noise (Rahnev et al., 2011; Solovey, Graney, & Lau, 2015); that is, participants were more likely to say that items were presented in peripheral or unattended locations, even when they were not. These results were interpreted to reflect a subjective sense of inflated phenomenology in the unattended periphery, because detection biases could in principle reflect both subjective perception and decisional or response strategies (Witt, Taylor, Sugovic, & Wixted, 2015). It was argued that a decisional or cognitive account of these biases is less likely because results remain consistent even when subjects are given feedback and training (Rahnev et al., 2011; Solovey et al., 2015).

If liberal biases reflect inflated visual phenomenology, we should expect these results to emerge in naturalistic settings, too. This question is important, because as noted by previous research, findings from artificial tasks may not be representative of the perceptual and decisional strategies used by the brain in more ecologically valid settings (Felsen & Dan, 2005). Additionally, research indicates that while most processing of real-world scene information is intact with diminished attention, the depth of processing of this information does depend on attention (Groen, Ghebreab, Lamme, & Scholte, 2016). Therefore, results from attentional cueing tasks based on a few simple stimuli on a gray background may not generalize well to naturalistic settings where the visual scene is complex. To address the question of whether liberal detection biases are evident when making judgments in the unattended periphery, we conducted a study where subjects engaged in a naturalistic task of simulated driving while they looked for (i.e., detected) specific colors of pedestrians’ clothing in the visual periphery. Unlike in previous studies (Rahnev et al., 2011; Solovey et al., 2015), the stimuli were not degraded by noise or reduced contrast; instead, near-threshold task difficulty was created more naturalistically by having the pedestrians move at a fast speed. We reasoned that if the previous results (Rahnev et al., 2011; Solovey et al., 2015) were not due to idiosyncratic strategies adopted in artificial psychophysics experiments, here, subjects should exhibit similar liberal detection biases in the unattended periphery as well.

## Methods

### Participants

Twenty-six individuals (fifteen female, eleven male) participated in Experiment 1; mean age = 20.8, SD = 3.0. Fifteen individuals (twelve female, three male) participated in Experiment 2; mean age = 21.8, SD = 5.5. We conducted a power analysis based on previous research (i.e., Rahnev et al. 2011), and determined that the number of participants should be greater or equal to 15 (see Supplementary Methods). No statistical analyses reported in the paper were performed on partial data. All participants were students at the University of California-Los Angeles. All of them had normal or corrected-to-normal vision, were 18 years or older, and did not have a history of seizures, epilepsy, stroke, or head trauma. Each participant gave informed consent and received either credits or monetary compensation of $10 per hour. This study was approved by the UCLA Institutional Review Board.

### Stimuli and Apparatus

Stimuli were generated using Unreal Engine 4 (Epic Games, Cary, NC, USA) to create game-like naturalistic environments and stimuli. Participants were instructed to use four keys to accelerate (key ‘W’), brake (key ‘S’) and steer (key ‘A’ for turning left and key ‘D’ for turning right), similar to the conventions in many racing games. The overall experience of stimuli was similar to 3D-racing video games. Participants were instructed to drive a vehicle along the straight two-lane road in a small town (Fig. 1A) and stop at the stoplight at the first intersection. There was no other vehicle on the road. The vehicle was parked 82 meters (in virtual distance) away from the stop line at the intersection at the beginning of each trial. When the vehicle was 43 meters away from the stop line, in 80% of the trials, a pre-cue textbox was displayed for approximately four seconds in the upper center of the screen, reading either “Attention to the LEFT” or “Attention to the RIGHT” (Fig 1B). Participants were informed that if a pre-cue textbox appeared, there was 75% chance that the item they would have to answer a detection question about would be at the same location as indicated by the pre-cue (“valid” trials), and there was 25% chance that the item they would have to answer a detection question about would be at a different location as indicated by the pre-cue (“invalid” trials). They were instructed to allocate attention equally to both the left and right locations if no pre-cue text was presented (“baseline” trials), which accounted for 20% of the total trials.

**Figure 1.**
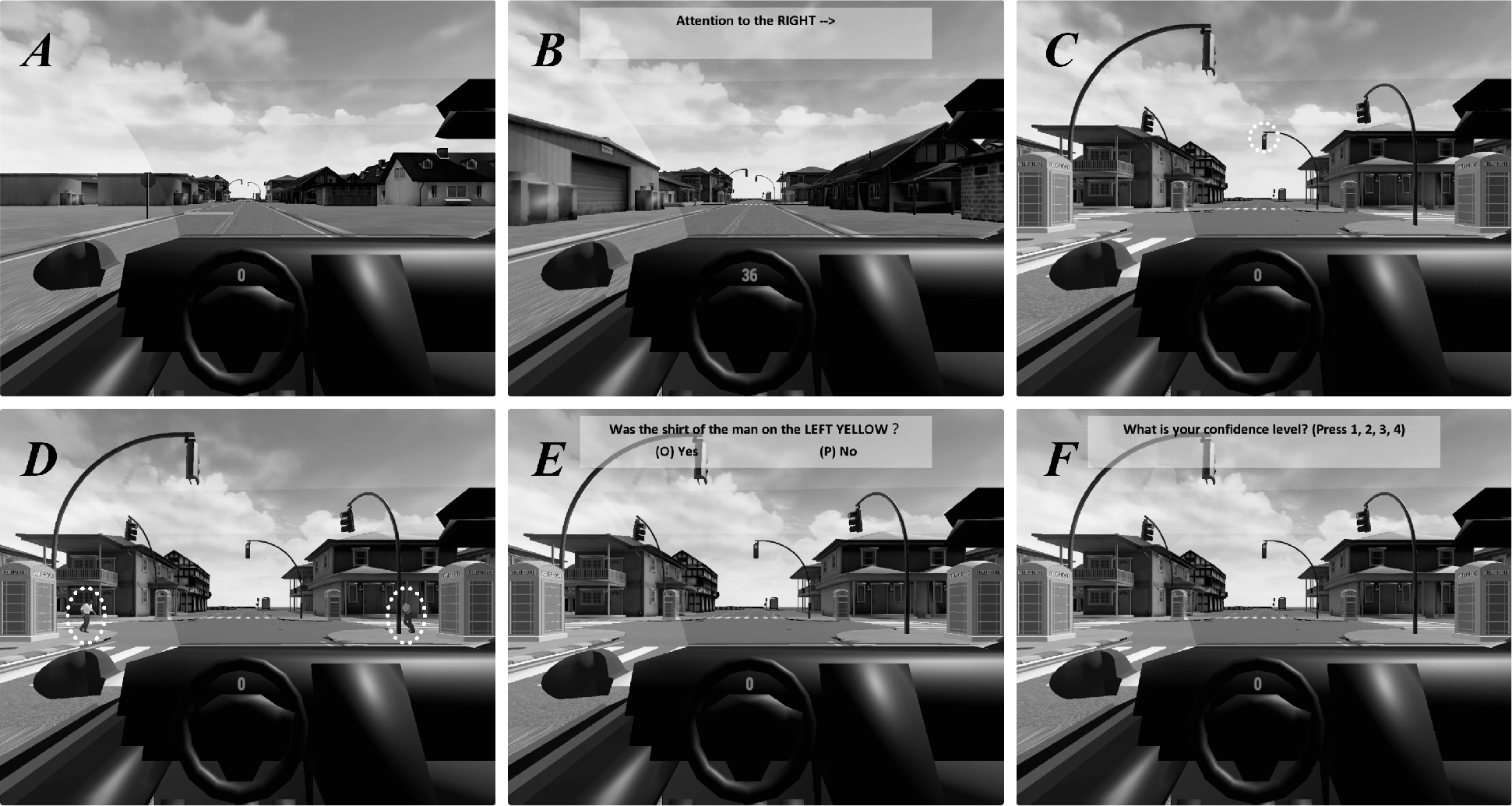
The task design of both experiments. (A) The start of each trial. Vehicles started from a parked position near the end of a straight, two-lane road in a small town, 82 meters away from the stop line of the intersection. (B) The attention cue. When the vehicle was 43 meters away from the intersection, in 80% of the trials, a pre-cue textbox was displayed in the upper central portion of the screen, which indicated the direction that subjects were to attend when stopped at the intersection. (C) Fixation validation. Before stimuli were presented, each participant’s fixation was continuously monitored to ensure he/she was fixating on the red stoplight for one full second. The white dotted circle indicates the position of the red light. (D) Presentation of stimuli. Two people wearing shirts of different colors ran out from behind two telephone booths positioned on each side of the street, and then turned around before getting halfway across the intersection. The two white, dotted circles indicate the positions of the two people. (E) The yes/no detection question. This question was presented one second after the stimuli disappeared. (F) The confidence question. After answering the yes/no question, participants were also asked to report their confidence level (1-4) regarding their choice.

Upon reaching the intersection, the vehicle was automatically moved to a fixed position to ensure the stimuli were presented in identical fashion on the screen on each trial. After the vehicle was fully stopped, the pre-cue text disappeared. Then, the participant’s fixation was continuously monitored to ensure the participant was fixating on the red light for one second (we set the tolerance of distance between fixation and center of the red light as 2.5^°^ in visual angle) (Fig. 1C). For details of eye tracking, see Supplementary Methods. The stimuli of interest in this task were two running male individuals, with shirt colors randomly drawn from seven potential colors for Experiment 1 (yellow, red, orange, green, blue, purple and gray) or eleven potential colors for Experiment 2 (red, pink, orange, yellow, green, blue, purple, brown, white, gray, and black) on each trial (Fig. 1D). These male pedestrians quickly ran out from behind telephone booths simultaneously at the speed of 17.5^°^ / second after the vehicle had stopped for one second, and then quickly ran back behind the telephone booths. They were presented for 300 milliseconds in total. 1000ms after the stimuli disappeared, a yes-no question (e.g., “Was the shirt color of the person on the left yellow?” or “Was the shirt color of the person on the right yellow?”) was presented (Fig. 1E). The probability that the correct choice was “Yes” or “No” were both 50%, and participants were told this explicitly in advance. In Experiment 1, the target color was always yellow, while in Experiment 2, the target color was randomly picked from the eleven possible colors in each trial. After answering the yes-no question, participants were also asked to report their confidence level regarding their choice from 1 (low confidence) to 4 (high confidence) (Fig. 1F).

Stimuli were shown on a 24-inch monitor (about 19-inch wide) with 60 Hz refresh rate and resolution of 1280 x 1024 pixels. Each shirt was approximately 1.1^°^ x 1.0^°^ in visual angles, and was located 15.4^°^ - 17.6^°^ away from fixation while the stimuli were moving. Participants were seated 60 cm from the screen, and viewed the screen freely with their head unrestrained.

### Procedure

In both experiments, all participants were required to complete two sessions; the two sessions occurred approximately one week apart. The first session included eye tracker calibration, 15 practice trials, and 210 experiment trials. The second session included eye tracker calibration and 270 experiment trials. Participants were instructed to take a break of up to 60 seconds after every 30 trials.

## Results

### Experiment 1: Evaluating detection biases for one specific color

To ensure participants were fixating as instructed, we analyzed their eye gaze at 17ms (i.e., time of one frame) before stimuli were presented, and during 300ms of stimuli presentation. Participants’ fixations when stimuli were presented from every trial in Experiment 1 are illustrated in Fig. 2A, which shows that subjects were fixating properly when stimuli were presented. The range of the target eccentricity at the onset of the stimulus is 15.52^°^ - 20.28^°^; the 99% confidence interval of the target eccentricity is (16.53^°^, 19.22^°^). Participants’ fixations during stimuli presentation are illustrated in Fig. 2B, which shows that the stimuli were presented in the periphery of our participants’ visual fields in nearly all of the trials. Trials in which participants directed their gaze towards the target were discarded, and are not included in the analyses in this manuscript.

**Figure 2.**
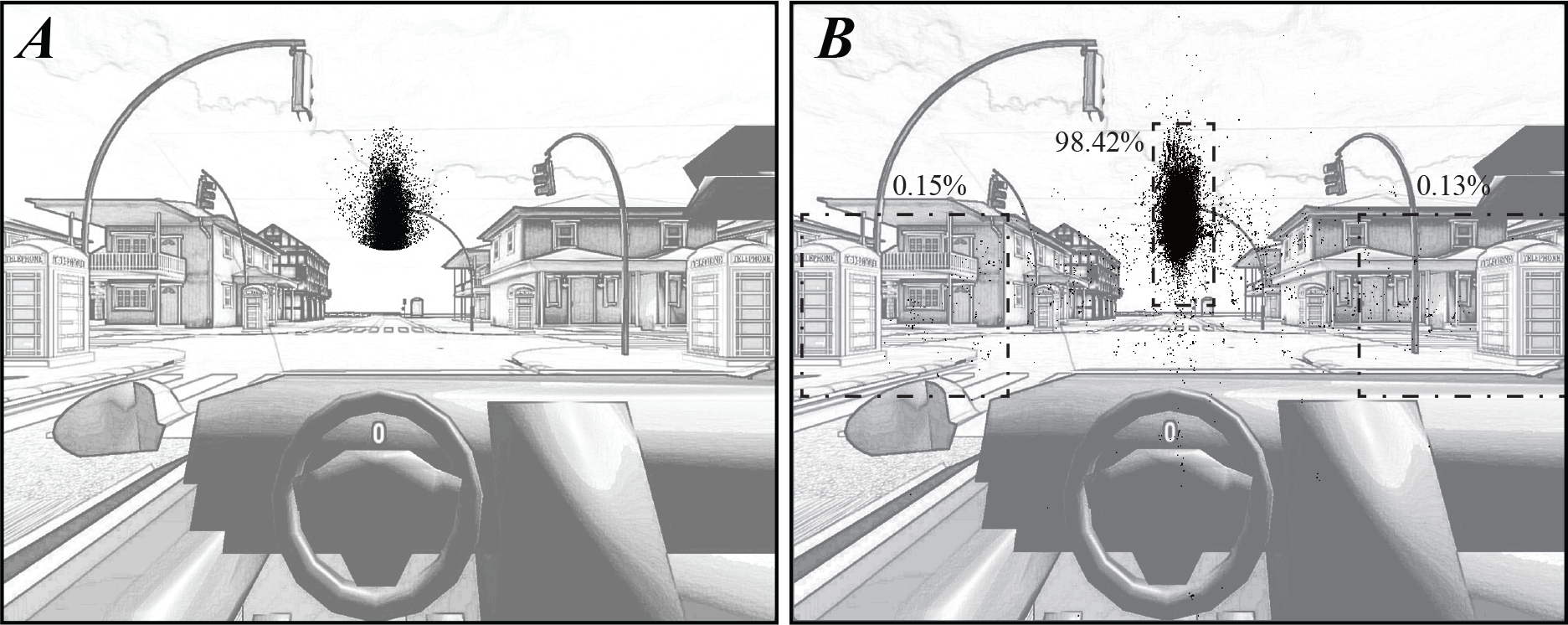
Participants’ fixations in Experiment 1. (A) Participants’ fixations at 17ms (i.e., time of one frame) before stimuli (i.e., the two male pedestrians) were presented, from every trial in Experiment 1. Each black point represents a fixation from one of the two eyes, and fixation points from both eyes are plotted in the figure. Almost all fixations are on or near the stoplight in every trial. (B) Participants’ fixations during stimulus presentation (300ms total) in every trial in Experiment 1. Each black point represents an average fixation of two eyes. Only about 0.28% of fixations are directed to the stimuli of interest (as located in black dash-dotted frames on the sides of the screen), while 98.42% of fixations were properly located near the traffic light (the black dashed frame in the middle).

Next, before conducting Signal Detection Theoretical (SDT) analyses, we assessed the equal variance assumption from SDT and found significant deviations in some subjects (Supplementary Methods, Supplementary Fig. 1). Because of this, in subsequent SDT analyses we used the measures *d_a_* (sensitivity) and *c_a_* (i.e., the criterion, or response bias) to account for unequal variances; this was possible because in addition to Yes/No answers, we also collected confidence ratings and could perform a full ROC analysis with multiple criteria points in ROC space for each subject.

For our primary analyses, we conducted Mauchly’s test and repeated measures ANOVAs to analyze the effects of attention on the percentage of correct responses, *d_a_* and *c_a_*. Overall, performance was similar across attention conditions (Fig. 3A), and the percentage of correct trials did not significantly differ across attention levels, *F*(2, 50) = 0.97, *p* = .387. This indicates the effects of attentional cuing were likely modest. However, performance was still highest in the valid attention trials and lowest in the invalid attention trials.

**Figure 3.**
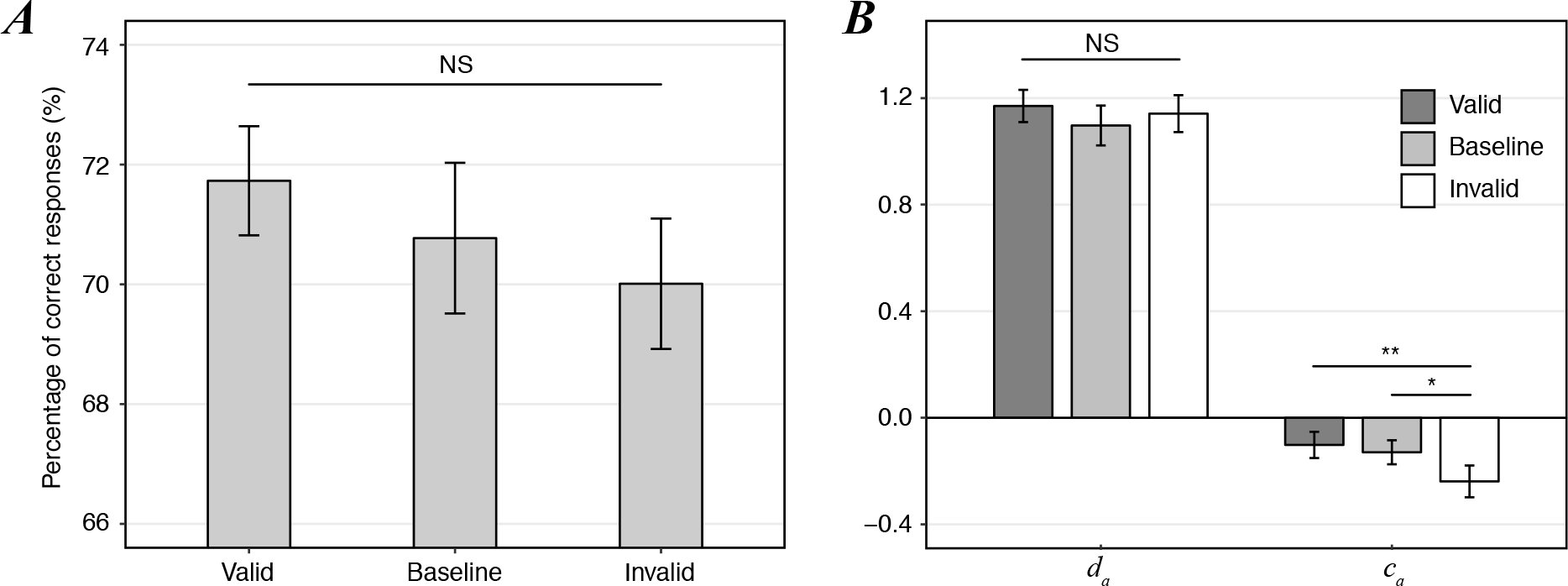
Behavioral results from Experiment 1. (A) The percentage of correct responses across attention conditions. While subjects exhibited the best performance for valid attention trials and the lowest performance for invalid trials, the conditions were not significantly different from one another. (B) *d_a_* and *c_a_* across attention conditions. *d_a_* values were quite consistent across attention conditions. *c_a_* under inattention (i.e., invalid trials) was significantly lower than *c_a_* in the valid or baseline conditions, providing evidence for a liberal detection bias. Bars represent the average values across subjects; error bars represent the standard error of the mean. ^**^*p* < .01, ^*^*p* < .05, NS: not significant.

Similar to the effect of attention on the percentage correct, the average *d_a_* was quite consistent across attention conditions (Fig. 3B) and *d_a_* did not significantly differ across attention levels, *F*(2, 50) = 0.43, *p* = .65. However, when we compared *c_a_* across attention conditions, *c_a_* values were significantly different across attention levels, *F*(2, 50) = 7.16, *p* = .002. Post-hoc paired *t*-tests indicated that the difference of *c_a_* between the valid condition and invalid condition was significant, *t*(25) = 3.20, *p* = .004; the difference of *c_a_* between baseline condition and invalid condition was also significant, *t*(25) = 2.70, *p* = .012; however, the difference of *c_a_* between valid condition and baseline condition was not significant, *t*(25) = 0.91, *p* = .371. This provided evidence that participants adopted more liberal criteria for making detection judgments when the target was unattended and presented in periphery.

### Experiment 2: Evaluating detection biases in the periphery for an array of colors

In Experiment 1, the detection question that was asked on every trial involved a constant target: a person in a yellow shirt. One could argue that a liberal detection bias may be observed due to the sheer fact that subjects may have a tendency to respond “Yes” whenever they are uncertain. That is, our primary finding of a liberal detection criterion could be driven by a confirmation bias at the decisional level. To test if this is the case, we designed Experiment 2 in a specific manner: we changed the target color on each trial, and participants only knew of the color they were to detect when they were prompted by the detection question at the end of the trial. This way, participants could not anticipate what target to detect beforehand. If the liberal detection bias we observed in Experiment 1 were due to a confirmation bias in responding, we would still see the effect here. On the other hand, we would not expect similar results if they were contingent on the fact that in Experiment 1 subjects were able to form a template for the target color before they were prompted.

As in Experiment 1, we evaluated participants’ fixations at 17ms before stimuli were presented and during stimuli presentation. The fixation positions indicated that stimuli were indeed presented in the periphery of our participants’ visual fields in nearly all trials (Supplementary Fig. 3). The range of the target eccentricity at the onset of the stimulus is 15.72^°^ - 21.05^°^; the 99% confidence interval of the target eccentricity is (16.40^°^, 19.67^°^). We discarded trials in which participants directed their gaze towards the target.

We evaluated the percent correct in each of the three attention conditions from Experiment 2, and also calculated *d_a_* and *c_a_* to measure the sensitivity and response criterion for each subject. Specifically, we conducted Mauchly’s test and repeated measures ANOVAs to evaluate the effect of attention on the percent correct, *d_a_* and *c_a_*. Overall, attention increased the percentage of correct responses, as participants exhibited the highest performance in the valid attention condition and the lowest performance in the invalid condition (Fig. 4A). The percentage of correct trials was significantly different across attention levels, *F* (2, 50) = 5.53, *p* = .009. Post-hoc paired *t*-tests indicated that the difference in the percentage of correct responses between the valid condition and invalid condition was significant, *t*(14) = 2.99, *p* = .01; the difference in the percentage of correct responses between the valid condition and baseline condition was also significant, *t*(14) = 2.39, *p* = .031, which suggested that participants performed better when allocating full attention to stimuli, rather than distributing attention across the two locations; the difference in the percentage of correct responses between the valid condition and baseline condition was not significant, *t*(14) = 1.52, *p* = .15.

**Figure 4.**
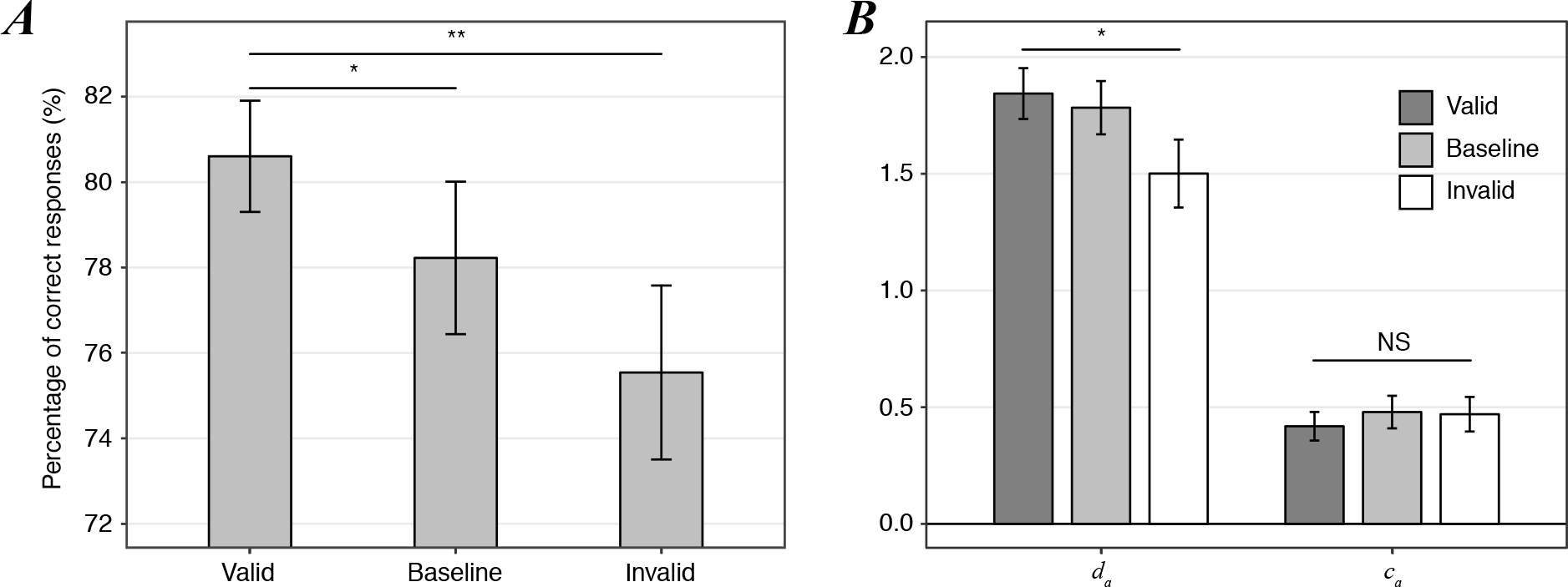
Behavioral results for Experiment 2. (A) The percentage of correct responses across attention conditions. The percentage of correct responses for valid trials was significantly larger than the percentage correct for baseline and invalid trials. (B) *d_a_* and *c_a_* across attention conditions. Similar to the effect of attention on percentage of correct responses, *d_a_* was significantly greater in the valid attention condition compared to the invalid condition. *c_a_* did not significantly differ across attention conditions, which indicated that participants used similar internal criteria to make perceptual judgments in all attention conditions. Bars represent means, error bars represent SEM. \**p* < .05, NS: not significant.

The average *d_a_* showed a similar trend compared to the percentage of correct responses: *d_a_* increased with attention significantly (Fig. 4B), *F* (1.45, 20.28) = 4.54, *p* = .033 (the degrees of freedom were corrected using Greenhouse-Geisser estimates of sphericity, *ε* = .72). Post-hoc paired *t*-tests indicated that the difference of *d_a_* between the valid condition and invalid condition was significant, *t*(14) = 2.46, *p* = .028, while the difference of *d_a_* between the baseline condition and invalid condition was not significant, *t*(14) = 2.04, *p* = .06; the difference of *d_a_* between the valid condition and baseline condition was also not significant, *t*(14) = 0.79, *p* = .44. As for the response criteria *c_a_*, the results showed that *c_a_* did not significantly differ across attention levels, *F*(2, 28) = 0.52, *p* = .6. Importantly, these response criteria were positive, meaning they were much more conservative than those observed in Experiment 1. This finding indicates that our results in Experiment 1 were not due to a sheer confirmation bias at the decisional level (see Discussion).

## Discussion

In this study, we investigated how subjects detected colorful stimuli in the unattended periphery in a naturalistic environment. In our first experiment, on each trial, we asked observers whether an individual wearing a shirt with a specific color (yellow) had been presented at a specific location. Results showed that observers exhibited higher numbers of false alarms (Supplementary Fig. 4) (i.e., saying a yellow shirt was present, even when it was not) in unattended locations in the visual periphery, compared to locations that were fully or partially attended. This tendency to use liberal detection criteria in unattended or peripheral locations has been shown previously in artificial settings (Rahnev et al., 2011; Solovey et al., 2015). Here, we confirmed the hypothesis that this detection bias extends to more naturalistic stimuli and tasks, too. In our second experiment, on each trial, we asked observers whether an individual wearing a shirt with a particular color had been presented at a specific location, but in this experiment, we varied the target color randomly from trial to trial. Results showed that subjects used relatively conservative perceptual criteria (i.e., were relatively reluctant to say a color was present) when making detection-related judgments in this experiment, regardless of the amount of attention that was allocated to a given location.

One may notice that observers showed dramatic differences in the placement of the perceptual criteria between our two experiments. We can explain the differences based on variance reduction model (Rahnev et al., 2011; Solovey et al., 2015). This model assumes a single unified criterion across attention levels, and proposes reduced trial-by-trial variability of the internal perceptual signal when attended. Thus, the criterion is more liberal when unattended, which is what we have found in Experiment 1. However, this model works based on an a priori, well-defined stimulus dimension on which the subject can place the criterion to do the detection. If the feature to be detected can only be known after the presentation of stimulus (which was the major change in Experiment 2), one is not able to place the criterion in the same location for both the cued and uncued stimuli over many trials. As such, we would not predict the inflation effect in Experiment 2. and this is exactly what we obtained. The conservative criteria obtained in Experiment 2 further confirms that the effect of attention on detection bias is unlikely just due to a confirmation bias at the decisional level, i.e., a general tendency to answer “yes” to any question, when uncertain. This means the variance reduction model, which a priori can only apply in Experiment 1 but not Experiment 2, is an effective account of the differences in perceptual decision making between these two experiments.

One important question which remains is what it means when people say they see the target more often. Traditionally, it is thought that much of peripheral vision is ‘filled in’ via top-down mechanisms (Komatsu, 2006). However, it has also been reported that people trust unreliable, filled-in percepts more than percepts based on external input (Ehinger, Häusser, Ossandón, & König, 2017), suggesting that filling-in may not be the complete mechanism to explain peripheral phenomenology, and that decisional or metacognitive mechanisms are also involved. In our study, perhaps the findings of detection bias in the unattended periphery can also be interpreted as congruent with this account involving mechanisms at the decisional or metacognitive level. Importantly, just because the effect is to be thought of at the ‘decisional’ level does not mean that this is unrelated to perceptual phenomenology; criterion effects can also reflect subjective percept (Phillips, 2016; Witt et al., 2015). This interpretation is in line with previous findings that people tend to overestimate their ability to detect changes in change blindness experiments (Levin, Momen, Drivdahl, & Simons, 2000), and in a sense, people are not fully aware of their decreased capacity for color perception in the periphery (Cohen et al., 2016).

In terms of practical implications, if people tend to confirm what they expect whenever they don’t attend, this raises an important issue regarding driving safety. Many people are optimistic and tend to expect things to be positive (Sharot, 2011; Sharot, Guitart-Masip, Korn, Chowdhury, & Dolan, 2012; Sharot, Riccardi, Raio, & Phelps, 2007), so they may tend to mistakenly detect hazards that present in unattended periphery as no danger, while what is actually happening is they just don’t see the hazards. Similar results have been found in previous studies where drivers showed conservative criteria in hazard detection regardless of driving experience (Ventsislavova et al., 2016; Wallis & Horswill, 2007). Hazard detection is a special detection task, because the penalty for a miss and a false alarm is different: a miss may cause a crash, while a false alarm may only lead to unnecessary brakes; thus, the criterion in hazard detection task is closely related with driving safety. In our research, the task is to detect pedestrian when the vehicle has stopped. However, in real driving, most of the hazards are presented while driving, which is quite different from our task. Future research should more systematically address how inattention affects hazard detection judgment while driving. Specifically, while the impact of distractions (e.g., due to phone calls and text messaging) on driving performance is well known (Haigney, Taylor, & Westerman, 2000; Rumschlag et al., 2015), the impact on specific aspects of the perceptual decision making process remains relatively unexplored.

One limitation of our study is that the attention manipulation wasn’t overly strong in Experiment 1. Yet there is still a trend that the performance increased with attention, and we obtained the effect that people showed liberal detection bias in the unattended periphery. Also, in our study we compared the criteria when the naturalistic stimuli were presented in the periphery at equal eccentricity, between validly cued and invalidly cued conditions. It is possible that a stronger difference in criteria can be found when naturalistic stimuli are presented in the center vs. the periphery, as has been done in previous studies using artificial stimuli (Solovey et al., 2015). We hypothesize that the decisional effects we obtained here may generalize to these other conditions too, using naturalistic stimuli. We hope this can be tested in the future.

## Acknowledgments

This work was supported by a grant from the Air Force Office of Scientific Research (FA-9550-15-1-0110) to HL.

## References

Block, N. (2007). Consciousness, accessibility, and the mesh between psychology and neuroscience. The Behavioral and brain sciences, 30 (5-6), 481–99; discussion 499-548.

Block, N. (2011). Perceptual consciousness overflows cognitive access. Trends in cognitive sciences, 15 (12), 567–575.

Cohen, M. A., Dennett, D. C., & Kanwisher, N. (2016). What is the bandwidth of perceptual experience? Trends in cognitive sciences, 20 (5), 324–335.

Ehinger, B. V., Häusser, K., Ossandón, J. P., & König, P. (2017). Humans treat unreliable filled-in percepts as more real than veridical ones. eLife, 6.

Felsen, G., & Dan, Y. (2005). A natural approach to studying vision. Nature neuroscience, 8 (12), 1643–1646.

Grimes, J. A. (1996). On the failure to detect changes in scenes across saccades. In K. Akins (Ed.), Perception. Oxford University Press.

Groen, I. I. A., Ghebreab, S., Lamme, V. A. F., & Scholte, H. S. (2016). The time course of natural scene perception with reduced attention. Journal of neurophysiology, 115 (2), 931–946.

Haigney, D. E., Taylor, R. G., & Westerman, S. J. (2000). Concurrent mobile (cellular) phone use and driving performance: task demand characteristics and compensatory processes. Transportation research. Part F, Traffic psychology and behaviour, 3 (3), 113–121.

Komatsu, H. (2006). The neural mechanisms of perceptual filling-in. Nature reviews Neuroscience, 7 (3), 220–231.

Lamme, V. A. F. (2004). Separate neural definitions of visual consciousness and visual attention; a case for phenomenal awareness. Neural networks: the official journal of the International Neural Network Society, 17 (5-6), 861–872.

Lamme, V. A. F. (2006). Towards a true neural stance on consciousness. Trends in cognitive sciences, 10 (11), 494–501.

Levin, D. T., Momen, N., Drivdahl, S. B., & Simons, D. J. (2000). Change blindness blindness: The metacognitive error of overestimating change-detection ability. Visual cognition, 7 (1-3), 397–412.

Phillips, I. (2016). Naïve realism and the science of (some) illusions. Philosophical Topics, 44 (2), 353–380.

Rahnev, D., Maniscalco, B., Graves, T., Huang, E., de Lange, F. P., & Lau, H. (2011). Attention induces conservative subjective biases in visual perception. Nature neuroscience, 14 (12), 1513–1515.

Rensink, R. A., O’Regan, J. K., & Clark, J. J. (1997). To see or not to see: The need for attention to perceive changes in scenes. Psychological science, 8 (5), 368–373.

Rumschlag, G., Palumbo, T., Martin, A., Head, D., George, R., & Commissaris, R. L. (2015). The effects of texting on driving performance in a driving simulator: the influence of driver age. Accident analysis and prevention, 74, 145–149.

Sharot, T. (2011). The optimism bias. Current biology: CB, 21 (23), R941–5.

Sharot, T., Guitart-Masip, M., Korn, C. W., Chowdhury, R., & Dolan, R. J. (2012). How dopamine enhances an optimism bias in humans. Current biology: CB, 22 (16), 1477–1481.

Sharot, T., Riccardi, A. M., Raio, C. M., & Phelps, E. A. (2007). Neural mechanisms mediating optimism bias. Nature, 450 (7166), 102–105.

Simons, D. J., & Chabris, C. F. (1999). Gorillas in our midst: sustained inattentional blindness for dynamic events. Perception, 28 (9), 1059–1074.

Solovey, G., Graney, G. G., & Lau, H. (2015). A decisional account of subjective inflation of visual perception at the periphery. Attention, perception & psychophysics, 77 (1), 258–271.

Sperling, G. (1960). the Information Available in Brief Visual Presentations. Psychological Monographs: General and Applied, 74 (11), 1–29.

Ventsislavova, P., Gugliotta, A., Peña-Suarez, E., Garcia-Fernandez, P., Eisman, E., Crundall, D., & Castro, C. (2016). What happens when drivers face hazards on the road? Accident analysis and prevention, 91, 43–54.

Wallis, T. S. A., & Horswill, M. S. (2007). Using fuzzy signal detection theory to determine why experienced and trained drivers respond faster than novices in a hazard perception test. Accident analysis and prevention, 39 (6), 1177–1185.

Ward, E. J., Bear, A., & Scholl, B. J. (2016). Can you perceive ensembles without perceiving individuals?: The role of statistical perception in determining whether awareness overflows access. Cognition, 152, 78–86.

Witt, J. K., Taylor, J. E. T., Sugovic, M., & Wixted, J. T. (2015). Signal detection measures cannot distinguish perceptual biases from response biases. Perception, 44 (3), 289–300.

